# Comparative linkage mapping uncovers massive chromosomal inversions that suppress recombination between locally adapted fish populations

**DOI:** 10.1101/2021.10.18.464892

**Authors:** Maria Akopyan, Anna Tigano, Arne Jacobs, Aryn P. Wilder, Hannes Baumann, Nina O. Therkildsen

**Author notes:** Corresponding author: Maria Akopyan.

## Abstract

The role of recombination in genome evolution has long been studied in theory, but until recently empirical investigations had been limited to a small number of model species. Here we compare the recombination landscape and genome collinearity between two populations of the Atlantic silverside (*Menidia menidia*), a small fish distributed across the steep latitudinal climate gradient of the North American Atlantic coast. Using ddRADseq, we constructed separate linkage maps for locally adapted populations from New York and Georgia and their inter-population lab cross. First, we used one of the linkage maps to improve the current silverside genome assembly by anchoring three large unplaced scaffolds to two chromosomes. Second, we estimated sex-specific recombination rates, finding 2.75-fold higher recombination rates in females than males—one of the most extreme examples of heterochiasmy in a fish. While recombination occurs relatively evenly across female chromosomes, it is restricted to only the terminal ends of male chromosomes. Furthermore, comparisons of female linkage maps revealed suppressed recombination along several massive chromosomal inversions spanning nearly 16% of the genome and segregating between locally adapted populations. Finally, we discerned significantly higher recombination rates across chromosomes in the northern population. In addition to providing valuable resources for ongoing evolutionary and comparative genomic studies, our findings represent a striking example of structural variation that impacts recombination between adaptively divergent populations, providing empirical support for theorized genomic mechanisms facilitating adaptation despite gene flow.

## Introduction

Recombination is a fundamental evolutionary mechanism that influences genetic variation and adaptive trajectories. The exchange of alleles onto different genetic backgrounds as a result of recombination can both facilitate and impede adaptive evolution (Tigano and Friesen 2016). It can promote adaptation by generating novel combinations of beneficial haplotypes (Felsenstein 1974), or by breaking up genetic associations to allow the purging of deleterious mutations from adaptive haplotypes (Muller 1964). Conversely, recombination can disrupt favorable allelic combinations, which in turn can reduce the fitness of a population (Smith 1978; Altenberg and Feldman 1987). Understanding the role of recombination in facilitating responses to selection has been the subject of extensive theoretical study (Felsenstein 1974; Otto and Barton 1997; Barton and Charlesworth 1998; Otto and Lenormand 2002), and a growing body of empirical evidence has demonstrated that recombination varies highly among taxa and can contribute to different patterns of genetic diversity and divergence across species (Dapper and Payseur 2017; Ritz *et al*. 2017; Stapley *et al*. 2017). This supports the notion that recombination plays a crucial role in genome evolution.

Studies in a wide range of species have shown that recombination also tends to vary across the genome both within and among chromosomes (Begun and Aquadro 1992; Wu *et al*. 2003; Anderson *et al*. 2006; Kim *et al*. 2007; Branca *et al*. 2011; Hinch *et al*. 2011; Haenel *et al*. 2018). In most cases, recombination is reduced at the center of chromosomes, with the rate of crossovers gradually increasing towards the telomeres. This variation in recombination along the genome – the recombination landscape – has a profound impact on the efficacy of selection. Genomic features that can alter recombination rates and maintain linkage between adapted alleles in the presence of gene flow may be favored by selection (Noor *et al*. 2001; Rieseberg 2001; Nosil *et al*. 2009). Structural rearrangements including inversions, translocations, and fusions, can thus have a considerable effect on genetic transmission by interfering with recombination and promoting genome divergence (Tigano and Friesen 2016; Wellenreuther and Bernatchez 2018).

Among structural variants, chromosomal inversions are known to strongly shape local recombination landscapes (Stevison *et al*. 2017). The key evolutionary effect of inversions is that they suppress recombination in a heterozygous state (Sturtevant and Beadle 1936). By suppressing recombination in heterokaryotypes, inverted chromosomal regions can capture multiple loci involved in adaptation to contrasting environments and protect these favorable combinations of adaptive alleles (Kirkpatrick and Barton 2006; Hoffmann and Rieseberg 2008; Yeaman 2013). Recombination continues normally in the homozygous state for inverted and uninverted haplotypes, respectively, allowing inversions to escape some of the deleterious consequences suffered when recombination is entirely suppressed (Kirkpatrick 2010). While inversion polymorphisms capturing locally adapted loci are predicted to be a favorable architecture for adaptation despite gene flow (Kirkpatrick and Barton 2006; Yeaman 2013), until recently, much of the evidence supporting the role of inversions in adaptation came from a few classic examples (Krimbas and Powell 1992; Stefansson *et al*. 2005; Joron *et al*. 2006).

Increasing accessibility to genomic sequence data has led to the discovery that structural genomic variants are associated with adaptive divergence in a wide range of species (Wellenreuther *et al*. 2019; Mérot 2020). For instance, inversions maintain genomic differentiation between migratory and stationary ecotypes of the Atlantic cod (*Gadhus morhua*; Kirubakaran *et al*. 2016; Sodeland *et al*. 2016). In the seaweed fly (*Coelopa frigida*), alternate haplotypes have opposing effects on larval survival and adult reproduction (Mérot *et al*. 2020). Clinal patterns of polymorphic inversions also underlie locally adapted ecotypes of a coastal marine snail (*Littorina saxatilis*; Faria *et al*. 2019a) and have played an important role in repeated evolution of marine and freshwater sticklebacks (*Gasterosteus aculeatus*; Jones *et al*. 2012; Roesti *et al*. 2015). While many examples of inversions associated with local adaptations come from aquatic systems where there is typically high gene flow counteracting adaptive divergence among populations, there is also evidence of chromosomal rearrangements facilitating adaptation to terrestrial environments (e.g., Christmas *et al*. 2019; Todesco *et al*. 2020; Hager *et al*. 2021). Despite a growing appreciation for the effects of recombination on the dynamics of selection, the genomic features affecting the recombination landscape are still poorly understood in many systems because most studies have historically been limited to inbred lines of cultivated or model species (Stapley *et al*. 2017). Even less is known about the variation in recombination rates and genome structure across diverging populations of the same species (Samuk *et al*. 2020; Schwarzkopf *et al*. 2020), especially in an ecological context—i.e., non-model natural populations examined across varying environments (Stapley *et al*. 2017).

Distributed across the world’s steepest latitudinal climate gradient along North America’s Atlantic coast (Baumann and Doherty 2013), Atlantic silversides (*Menidia menidia*, hereafter: silversides) exhibit a remarkable degree of local adaptation in a suite of physiological and morphological traits (Conover *et al*. 2005). For example, the species exhibits countergradient variation in growth capacity (Conover and Present 1990), whereby northernmost populations have evolved higher growth capacity in response to shorter growing seasons, whereas tradeoffs with predator avoidance have selected for slower growth in the south (Billerbeck *et al*. 2001; Munch and Conover 2003; Arnott *et al*. 2006). Silverside populations also exhibit clinal genetic variation in vertebral number, temperature-dependent sex determination, swimming performance, lipid storage, spawning temperature and duration, egg volume, egg production, and size of offspring at hatch (Conover *et al*. 2009). Due to their broad distribution, abundance, and relative ease of husbandry, Atlantic silversides have been the focus of a wide range of ecological and evolutionary studies, such as experiments on fisheries-induced evolution (Conover and Munch 2002), responses to climate change (DePasquale *et al*. 2015; Murray *et al*. 2016), and local adaptation (Conover and Heins 1987; Conover and Present 1990; Schultz *et al*. 1998). However, after decades of research, we are only just beginning to explore the genomic basis underlying the remarkable capacity for adaptation in this ecological and evolutionary model species (Therkildsen *et al*. 2019; Therkildsen and Baumann 2020; Wilder *et al*. 2020; Tigano *et al*. 2021a).

Our recent work started to examine the genomic basis of local adaptation in silversides, revealing variation in genome structure among populations. We discovered that despite high gene flow maintaining overall low levels of genome divergence between populations, large blocks of the genome show strong linkage disequilibrium (LD) and differentiation between populations (Wilder *et al*. 2020; Tigano *et al*. 2021a). Strong LD spanning millions of bases, including thousands of variants fixed for alternate alleles in different populations, supported the presence of chromosomal inversions that maintain divergent adaptive haplotypes between highly connected silverside populations (Therkildsen *et al*. 2019; Therkildsen and Baumann 2020; Wilder *et al*. 2020; Tigano *et al*. 2021a). Subsequent alignments of genome assemblies from northern and southern populations and comparative analysis of the linear order of scaffolds resolved with Hi-C data confirmed that these blocks of divergence indeed represent inversions (Tigano *et al*. 2021a). Examining how inversions, both in their homozygous and heterozygous states, impact recombination patterns across locally adapted populations is an important next step in understanding the genomic architecture of adaptation. Thanks to the availability of a chromosome-level reference genome and the ability to create lab crosses, we conducted comparative linkage mapping to describe the recombination landscapes of Atlantic silversides within two adaptively divergent populations and their inter-population cross. We first used these maps to anchor large unplaced scaffolds to our previously published silverside genome assembly. We then compared the ordering and genetic distance between markers in the different linkage maps to their physical positions in the genome assembly to identify chromosomal rearrangements and calculate recombination rates. These comparisons allowed us to examine how recombination rates vary across central vs. terminal and inverted vs. uninverted regions of different chromosomes and how recombination differed between sexes and populations.

## Methods

### Mapping families

We generated three crosses for linkage mapping, including two F1 families resulting from reciprocal crossing of wild-caught silversides from two adaptively divergent parts of the distribution range (Georgia and New York), and one F2 family from intercrossing lab-reared progeny from one of the F1 families (Figure 1). Because linkage mapping measures recombination during gamete production in the parents, the F1 families give us separate information about the wild-caught male and female founder fish from each separate population (the F0 progenitors), and the F2 map reflects recombination in the hybrid F1 progeny.

**Figure 1.**
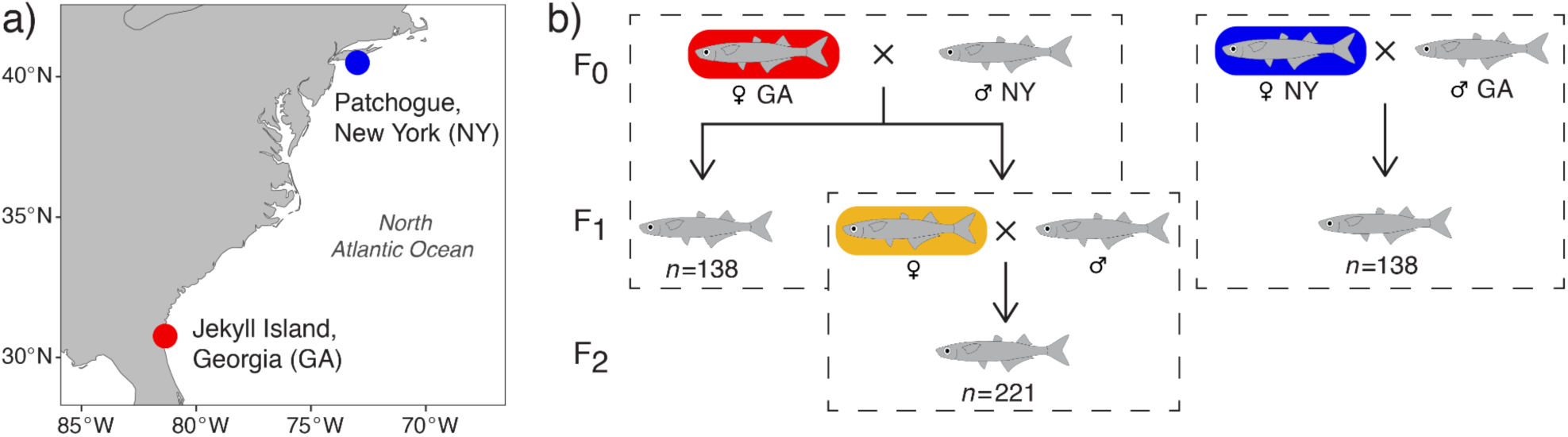
Experimental Design Map of sampling localities of wild-caught F0 individuals from Jekyll Island, Georgia and Patchogue, New York (a) used for generating mapping families (b). Each dashed box represents a family for which we produce a linkage map and the number of offspring (n) analyzed in each family is labeled. Focal females used for population comparisons are colored to match Figures 5-7 (and sampling localities in the founders).

In the spring of 2017, spawning ripe founders were caught by beach seine from Jekyll Island, Georgia (31°03’N, 81°26’W) and Patchogue, New York (40°45’N, 73°00’W) and transported live to the Rankin Seawater Facility at University of Connecticut’s Avery Point campus. For each family, we strip-spawned a single male and a single female onto mesh screens submerged with seawater in plastic dishes, then transferred the fertilized embryos to rearing containers (20 l) placed in large temperature-controlled water baths with salinity (30 psu) and photoperiod held constant (15L:9D). Water baths were kept at 20°C for the New York mother and at 26°C for Georgia mother families, which increased hatching success by mimicking the ambient spawning temperatures at the two different latitudes. Post hatch, larvae were provided *ad libitum* rations of newly hatched brine shrimp nauplii (*Artemia salina*, brineshrimpdirect.com). At 22 days post hatch (dph), we sampled 138 full-sib progeny from each of the two F1 families to be genotyped. The remaining offspring from the Georgia F1 family were reared to maturity in groups of equal density (40-50 individuals). In spring 2018, one pair of adult F1 siblings from the Georgia family were intercrossed to generate the F2 mapping population. At 70 dph, we sampled 221 full-sib F2 progeny for genotyping. In total, we analyzed 503 individuals: the two founders and 138 offspring from each of the two F1 families, plus two additional F1 siblings from the Georgia mother F1 family and their 221 F2 offspring (Fig. 1b). All animal care and euthanasia protocols were carried out in accordance with the University of Connecticut’s Institutional Animal Care and Use Committee (A17-043).

### Genotyping

We extracted DNA from each individual with a Qiagen DNeasy tissue kit following the manufacturer’s instructions and used double-digest restriction-site associated DNA (ddRAD) sequencing (Peterson *et al*. 2012) to identify and genotype single nucleotide polymorphisms (SNPs) for linkage map construction. We created two ddRAD libraries, each with a random subset of ∼250 barcoded individuals, using restriction enzymes *MspI* and *PstI* (New England BioLabs cat. R0106S and R3140S, respectively), following library construction steps as in Peterson *et al*. (2012). We size-selected libraries for 400-650 bp fragments with a Pippin Prep instrument (Sage Science) and sequenced the libraries across six Illumina NextSeq500 lanes (75 bp single-end reads) at the Cornell Biotechnology Resource Center.

Raw reads were processed in Stacks v2.53 (Catchen *et al*. 2013) with the module *process_radtags* to discard low-quality reads and reads with ambiguous barcodes or RAD cut sites. The reads that passed the quality filters were demultiplexed to individual fastq files. To capture genomic regions potentially not included in the current reference genome assembly, we ran the *ustacks* module to assemble RAD loci *de novo* (rather than mapping to the reference genome). We required a minimum of three raw reads to form a stack (i.e., minimum read depth, default -*m* option) and allowed a maximum of four mismatches between stacks to merge them into a putative locus (-*M* option).

Because the founders contain all the possible alleles that can occur in the progeny (except from any new mutations), we assembled a catalog of loci with *cstacks* using only the four wild-caught F0 progenitors. We built the catalog with both sets of founders to allow cross-referencing of common loci across the resulting F1 maps and we allowed for a maximum of four mismatches between loci (-*n* option). We matched loci from all progeny against the catalog with *sstacks*, transposed the data with *tsv2bam* to be organized by sample rather than locus, called variable sites across all individuals, and genotyped each individual at those sites with *gstacks* using the default SNP model (marukilow) with a genotype likelihood ratio test critical value (α) of 0.05.

Finally, we ran the *populations* module three times to generate a genotype output file for each mapping cross. For each run of *populations*, we specified the type of test cross (--*map-type* option cp or F2), pruned unshared SNPs to reduce haplotype-wise missing data (-*H* option), and exported loci present in at least 80% of individuals in that cross (-*r* option) to a VCF file, without restricting the number of SNPs retained per locus.

### Linkage mapping

We constructed separate linkage maps for each family using Lep-MAP3 (Rastas 2017), which can handle large SNP datasets and is appropriate for outbred families. For each map (made from a single set of parents and their offspring), we ran the SeparateChromosomes2 module to assign markers into linkage groups using segregation-distortion-aware logarithm of odds (LOD) scores (*distortionLod* = 1), following the author’s recommendations for single family data (Rastas 2018). We tested a range of 10 to 25 for LOD score thresholds (*lodLimit*) and evaluated the resulting number of linkage groups and the assignment distribution of markers to each linkage group. The LOD score thresholds were chosen based on variation in size between the largest linkage groups as well as the tail distribution of linkage group size (Rastas 2018). For each map, we chose the smallest LOD threshold at which the largest linkage groups were not further separated and increasing the threshold would instead add smaller linkage groups with few markers while the majority of markers remained in the largest groups.

Next, we used the *OrderMarkers2* module to order markers and compute genetic distances in centimorgan (i.e., recombination frequency, cM) between all adjacent markers for each linkage group using the default Haldane’s mapping function. We repeated this analysis for each parent in both F1 families: we used maternally informative markers (i.e., markers that were heterozygous only in the mother) to estimate recombination between alleles of the F0 female, and paternally informative markers (i.e., markers that were heterozygous only in the father) to estimate recombination between alleles of the F0 male. To investigate sex-specific heterogeneity in recombination, we calculated ratios between female and male total map distances for each linkage group. We then compared our findings to a recent metanalysis of sex-specific recombination rate estimates for 61 fish species (Cooney *et al*. 2021). We replicated their analysis by recalculating map length for males and females as the residuals of the relationship between log10-transformed map lengths and number of markers to control for the effect of marker number on map length estimates.

Due to strong heterochiasmy, i.e., different recombination rates between the sexes, with male recombination restricted to the terminal ends in most linkage groups (details below), we focused our cross-population comparisons on the female linkage maps in the remainder of the analyses. While we used only maternally informative markers to generate female maps from the two F1 families (Figure 1, red and blue), we used both maternally informative and dually informative markers to get comparable resolution (number of markers) for tracking segregation patterns in the F2 family hybrid mother (Figure 1, yellow). Depending on the genotypes of the F1 individuals sampled to generate the F2 family, a marker that was informative in one parent can (i) remain informative, (ii) can become dually informative if two heterozygous F1s were crossed, or (iii) can become uninformative if two homozygous F1s were crossed (Figure S1). As a result, in the F2 family the number of maternally and paternally informative markers is reduced, but some of SNPs that were uninformative in the F1s because the founders were homozygous for different alleles become dually informative for the F2 generation.

### Genome anchoring and improved assembly

We aligned the catalog of RAD loci to our recently published silverside reference genome (Tigano *et al*. 2021a) using Bowtie2 v2.2.9 (Langmead and Salzberg 2012) with the --*very*-*sensitive* preset option and converted alignments to BAM output with Samtools v1.11 (Li *et al*. 2009). Using the *Stacks* script *stacks_integrate_alignments*, we generated a table of genome coordinates for SNPs in the catalog BAM file, which we subsequently used to extract the physical positions of markers in the linkage maps. Although the reference genome is largely assembled to chromosome level with a scaffold N50 of 18.19 Mb and contains 89.6% BUSCO genes, a number of scaffolds remain unplaced. Therefore, we used the Georgia linkage map (as the reference genome was built with samples from this location) to aid placement of these scaffolds.

We anchored and reassembled the silverside genome with Lep-Anchor (Rastas 2020). To construct a chain file of contig–contig alignments, which Lep-Anchor takes as input data for linking genome contigs, we ran the first two steps of the HaploMerger2 (Huang *et al*. 2017) pipeline on the silverside genome with repeats masked using Red (Girgis 2015; Huang *et al*. 2017), a repeat-detection tool that applies machine learning to label training data and train itself automatically on an entire genome. In addition to the Georgia linkage map, the pairwise contig alignment data are used to infer the orientation and placement of contigs by maximizing the correlation of the physical (base pair) and the linkage map (cM) positions in Lep-Anchor’s *PlaceAndOrientContigs* module. We assigned previously unassembled genome scaffolds greater than 1 Mb (*n*=3, Tigano *et al*. 2021) to the 24 largest scaffolds in the reference genome to generate the linkage-map anchored assembly.

### Synteny analysis and recombination rate estimation

To examine how genetic distance and ordering between markers in the different linkage maps compares to the physical distance on the reassembled chromosomes, we constructed Marey maps that illustrate the position of each SNP in a linkage map against its coordinate in our anchored genome assembly. We initially included all SNPs per RAD-tag to maximize the number of informative markers for linkage mapping, then filtered to retain only one SNP per RAD-tag to reduce redundant data in subsequent analyses. We also removed a small number of outlier SNPs in the Marey map of each chromosome that disrupted the monotonically increasing trend expected from a Marey map function, as these can represent errors in the genetic and/or physical map (Marey maps including the outliers are shown in Figures. S5-S7). By comparing genetic positions from each linkage map to the physical positions in the linkage-map anchored assembly, we identified chromosomal rearrangements as regions containing more than 10 markers with a trend deviating from the linear alignment. We approximated inversion breakpoint locations as the mid-point between the physical coordinates of the markers flanking the edges of identified inverted regions.

To estimate broad-scale variation in recombination rates for each linkage group in each of the three female maps, we divided the length of the linkage group in cM by the length of the scaffold in Mb. To compare recombination rates of the three female maps while accounting for chromosome size, we ran an ANCOVA followed by post hoc analysis with a Bonferroni adjustment. In addition, we used the BREC package (Mansour *et al*. 2021) for estimating local recombination rates in each of the three maps. First, we used the filtered Marey map data to reverse the marker order of regions that are inverted compared to the linkage-map anchored assembly for each of the three female linkage maps. Then, we estimated local recombination rates using the Marey map approach with the linearized markers by correlating genetic and physical maps and fitting a local regression model (Loess with span 0.15).

To compare how fine-scale recombination rates vary between and within chromosomes with and without inversions, we analyzed recombination rates (from the Loess model) using a linear mixed model fit by maximum likelihood with two fixed factors: chromosomal region and mapping family, then used least-square means for post-hoc pairwise comparisons. For this analysis, we compared chromosomes that are collinear among the three female maps (i.e., no inversions) to chromosomes with alternate inversion arrangements in Georgia and New York. We further classified chromosomes into terminal regions (20% of physical length made up of 10% from each end), inverted regions (for chromosomes with inversions), and central regions (not terminal and outside inversions). Statistical analyses were conducted in R v. 3.6.1 (Team 2020) using package lme4 (Bates *et al*. 2011).

## Results

### Genotyping and linkage map construction

We obtained 1,840,133,831 raw reads from the 503 silverside samples with an average of 3,658,318 reads per sample. After adapter trimming and quality filtering, we retained 1,709,540,728 reads (93%), with an average of 3,398,689 reads per sample. We identified 236,608 loci across all samples, with an average of 45.7% of loci present in each sample (stdev=3.9%, min=0.007%, max=63%), and 19.1x mean per-sample coverage for loci present in the sample (stdev=4.1x, min=6.2x, max=31.2x). Following genotyping and filtering (>80% individuals genotyped per family), we retained 60,671 SNPS across 54,937 loci in the Georgia mother F1 family, 64,389 SNPs across 56,028 loci in the New York mother F1 family, and 59,926 variant sites across 54,526 loci in the F2 family.

Only a subset of the identified SNPs are informative for linkage map construction since linkage can only be determined between markers in which the focal parent has a heterozygous genotype. We were able to use 18,285 female informative and 19,820 male informative markers in the Georgia mother F1 family, 20,240 female informative and 19,662 male informative markers in the New York mother F1 family, and 20,696 female and dually informative markers in the F2 family. In each of the genetic maps, we obtained 24 linkage groups, consistent both with the haploid number of *M. menidia* chromosomes inferred from karyotyping (Warkentine *et al*. 1987) and the number of putative chromosome clusters identified in both populations with Hi-C data (Tigano *et al*. 2021a). Broadly speaking, the linkage groups are relatively homogenous in the number of markers across all maps. The total lengths and the number of markers in linkage groups in each of the resulting maps are summarized in Table 1.

**Table 1.** Summary of the total lengths (in cM) and number of SNPs assigned to the different linkage groups (LG) in each of the male and female linkage maps. Downstream analyses that compare linkage map positions to the physical position of markers in the genome sequence are based only on the subset of markers shown here that map to the genome assembly, survived manual outlier removal, and include only one SNP per RAD locus (map lengths and SNP counts retained for analysis are shown in Table S1).

| LG | Female |  |  |  |  |  | Male |  |  |  |
| --- | --- | --- | --- | --- | --- | --- | --- | --- | --- | --- |
|  | Georgia |  | New York |  | F1 |  | Georgia |  | New York |  |
|  | cM | SNPs | cM | SNPs | cM | SNPs | cM | SNPs | cM | SNPs |
| 1 | 195.6 | 784 | 201.3 | 928 | 170.0 | 886 | 112.7 | 887 | 88.8 | 870 |
| 2 | 195.6 | 444 | 202.0 | 513 | 178.6 | 339 | 58.0 | 432 | 30.4 | 416 |
| 3 | 195.6 | 924 | 201.3 | 1033 | 128.3 | 1102 | 63.9 | 1024 | 69.9 | 1093 |
| 4 | 152.3 | 988 | 202.0 | 998 | 127.7 | 1190 | 68.8 | 995 | 54.9 | 1019 |
| 5 | 195.6 | 934 | 204.8 | 1038 | 178.6 | 1034 | 69.6 | 986 | 63.9 | 943 |
| 6 | 195.6 | 995 | 160.6 | 1093 | 177.2 | 887 | 72.1 | 1011 | 52.7 | 1064 |
| 7 | 194.9 | 1021 | 199.8 | 1020 | 176.4 | 1025 | 49.8 | 1043 | 54.6 | 1027 |
| 8 | 193.4 | 523 | 162.1 | 873 | 178.6 | 1109 | 57.6 | 1144 | 57.9 | 943 |
| 9 | 195.6 | 755 | 202.0 | 832 | 178.6 | 829 | 75.0 | 817 | 74.1 | 876 |
| 10 | 146.4 | 859 | 165.5 | 887 | 178.6 | 856 | 61.7 | 867 | 50.3 | 839 |
| 11 | 158.9 | 772 | 160.8 | 983 | 147.3 | 1061 | 49.9 | 1033 | 53.3 | 959 |
| 12 | 195.6 | 710 | 163.5 | 811 | 178.1 | 747 | 58.7 | 775 | 53.6 | 759 |
| 13 | 195.6 | 874 | 160.4 | 894 | 178.1 | 880 | 88.1 | 851 | 79.8 | 845 |
| 14 | 138.7 | 791 | 202.0 | 856 | 146.0 | 923 | 54.6 | 856 | 60.6 | 811 |
| 15 | 155.5 | 799 | 177.3 | 837 | 175.9 | 842 | 60.5 | 796 | 102.7 | 816 |
| 16 | 169.3 | 830 | 187.2 | 899 | 178.6 | 597 | 63.5 | 912 | 80.8 | 906 |
| 17 | 195.6 | 716 | 201.3 | 809 | 144.7 | 806 | 63.1 | 774 | 59.5 | 780 |
| 18 | 195.6 | 440 | 141.6 | 559 | 126.4 | 920 | 60.8 | 526 | 60.8 | 460 |
| 19 | 110.2 | 665 | 145.7 | 770 | 152.2 | 622 | 62.3 | 685 | 55.2 | 670 |
| 20 | 193.4 | 638 | 197.6 | 700 | 172.6 | 688 | 55.9 | 644 | 51.4 | 706 |
| 21 | 195.6 | 777 | 201.3 | 785 | 148.7 | 836 | 66.5 | 721 | 60.2 | 781 |
| 22 | 156.7 | 807 | 162.5 | 848 | 166.0 | 521 | 76.7 | 797 | 67.6 | 778 |
| 23 | 195.6 | 620 | 202.0 | 622 | 178.6 | 736 | 81.2 | 620 | 66.7 | 657 |
| 24 | 195.6 | 619 | 162.3 | 652 | 178.6 | 1260 | 116.1 | 624 | 61.3 | 644 |
| Total | 4313 | 18285 | 4366 | 20240 | 3944 | 20696 | 1647 | 19820 | 1511 | 19662 |

### Sex differences in recombination

Comparison of male and female linkage maps reveals conspicuous recombination suppression in males overall, with female maps on average 2.75 times longer than the male maps (Table 1). On all male chromosomes, recombination appears to be restricted to the terminal ends of each chromosome, in most cases to only one end of a chromosome, in both the Georgia male (Figure 2 and S8) and the New York male (Figure S9). Compared to the male and female map lengths for 61 fish species (Cooney *et al*. 2021), Atlantic silversides represent one of the most extreme examples of sex-biased recombination rates (Figure 3). When comparing raw map lengths, the highest female:male ratio (3.59:1) is reported for the Chinook salmon (*Oncorhynchus tshawytscha*; McKinney *et al*. 2016). However, this is partly attributable to the different number of markers in the male and female maps used in this study, and the signal is tempered when accounting for the difference in number of markers. In the zebrafish (*Danio rerio*), the ratio between female and male map lengths is 2.74 to 1 (Singer *et al*. 2002), only slightly lower than what we see in the Atlantic silverside. After transforming map lengths to account for different numbers of markers, the greatest difference in map lengths is seen in zebrafish and Atlantic silversides (Figure 3). In contrast to the male maps, the female maps show extensive recombination across the entire length of each chromosome, so we focus on the female maps for the analysis of synteny and recombination rate variation. Among the female linkage maps, we found the largest map length in the New York female (4366 cM) compared to the Georgia female (4313 cM) and the inter-population hybrid female (3944 cM).

**Figure 2.**
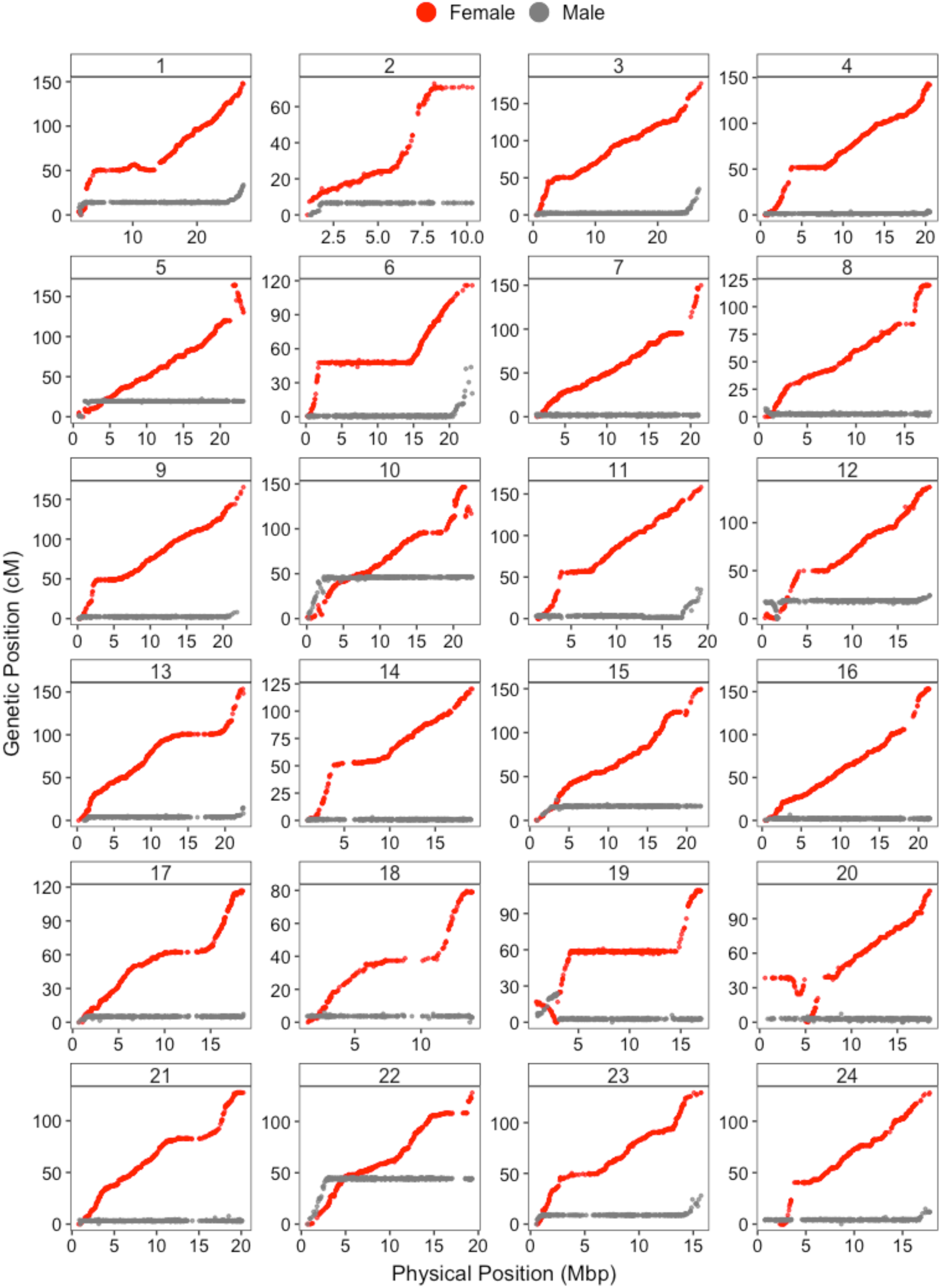
Male and Female Marey Maps The genetic map position (cM) vs. the physical position in the genome sequence of the SNPs assigned to each chromosome for the male and female from Georgia reveals extreme heterochiasmy, with male recombination restricted to the terminal ends in most linkage groups. These plots include only one SNP per RAD locus that maps to the reference genome (unfiltered male maps are shown in Figures S8 and S9).

**Figure 3.**
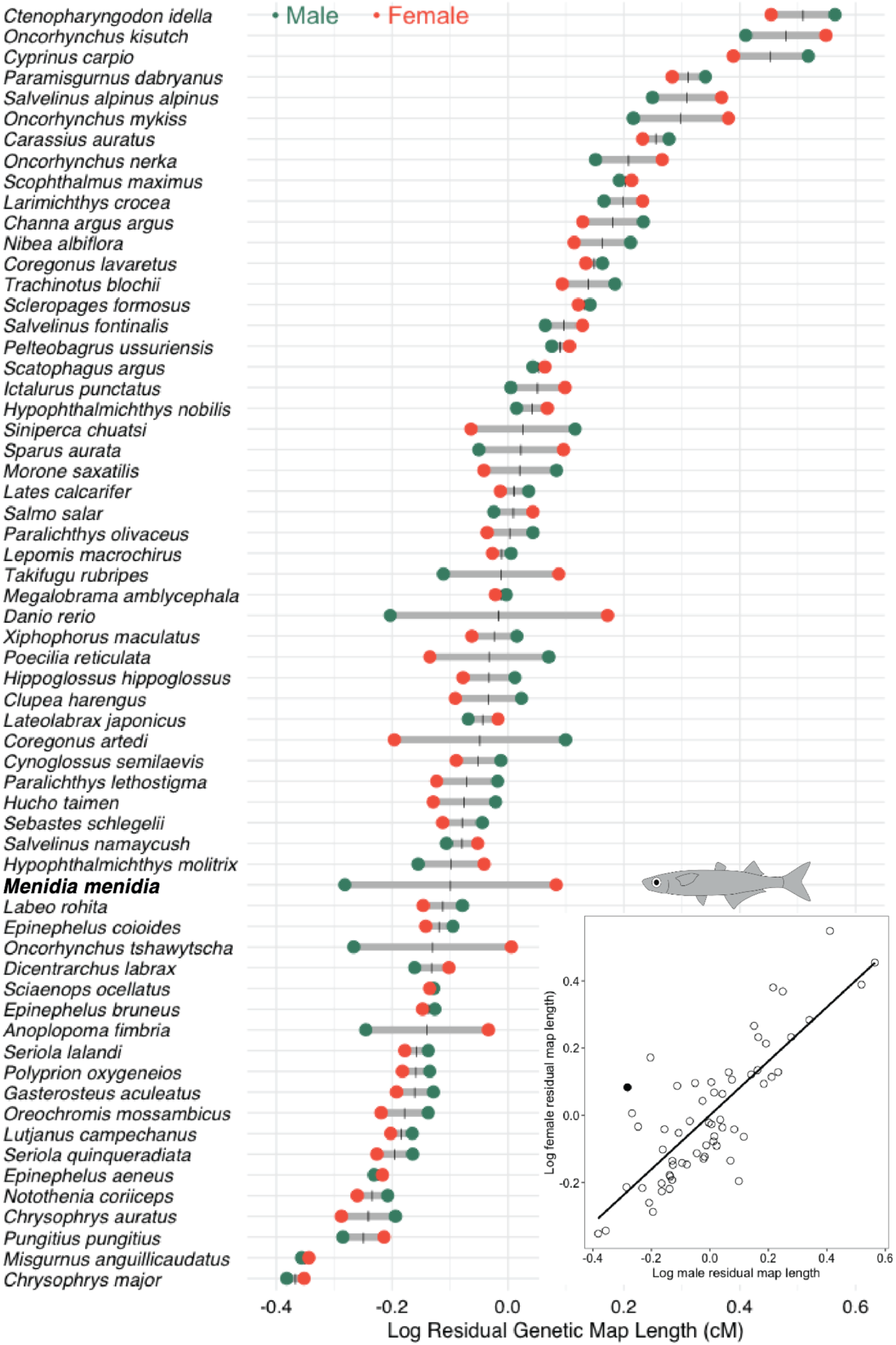
Heterochiasmy in Fishes Atlantic silversides have one of the highest differences between female and male map lengths compared to 61 fish species reviewed by Cooney *et al* 2021. For each species listed, map length after accounting for variation in numbers of markers is shown in green for males and orange for females. Grey horizontal bars represent the difference between the sexes and the vertical black bars indicate the sex-averaged map length, which was used to order species along the y-axis. The inset plot represents the relationship between male and female map lengths, where the Atlantic silverside (closed circle) shows considerable deviation from a one-to-one ratio.

### Linkage map anchored assembly

Considering only one SNP per RAD locus, 12,785 of the 18,285 SNPs in the Georgia female linkage map mapped to the reference genome and were used to anchor and order unplaced scaffolds into chromosome-scale pseudomolecules. Of these, 11,014 (86.1%) mapped to one of the main 24 scaffolds in the published genome assembly, 305 (2.4%) mapped to three additional scaffolds (>1 Mb), 830 (6.5%) mapped to smaller unplaced scaffolds, and 636 (5%) did not map to any sequence in the reference genome assembly (Figure 4 and S2).

**Figure 4.**
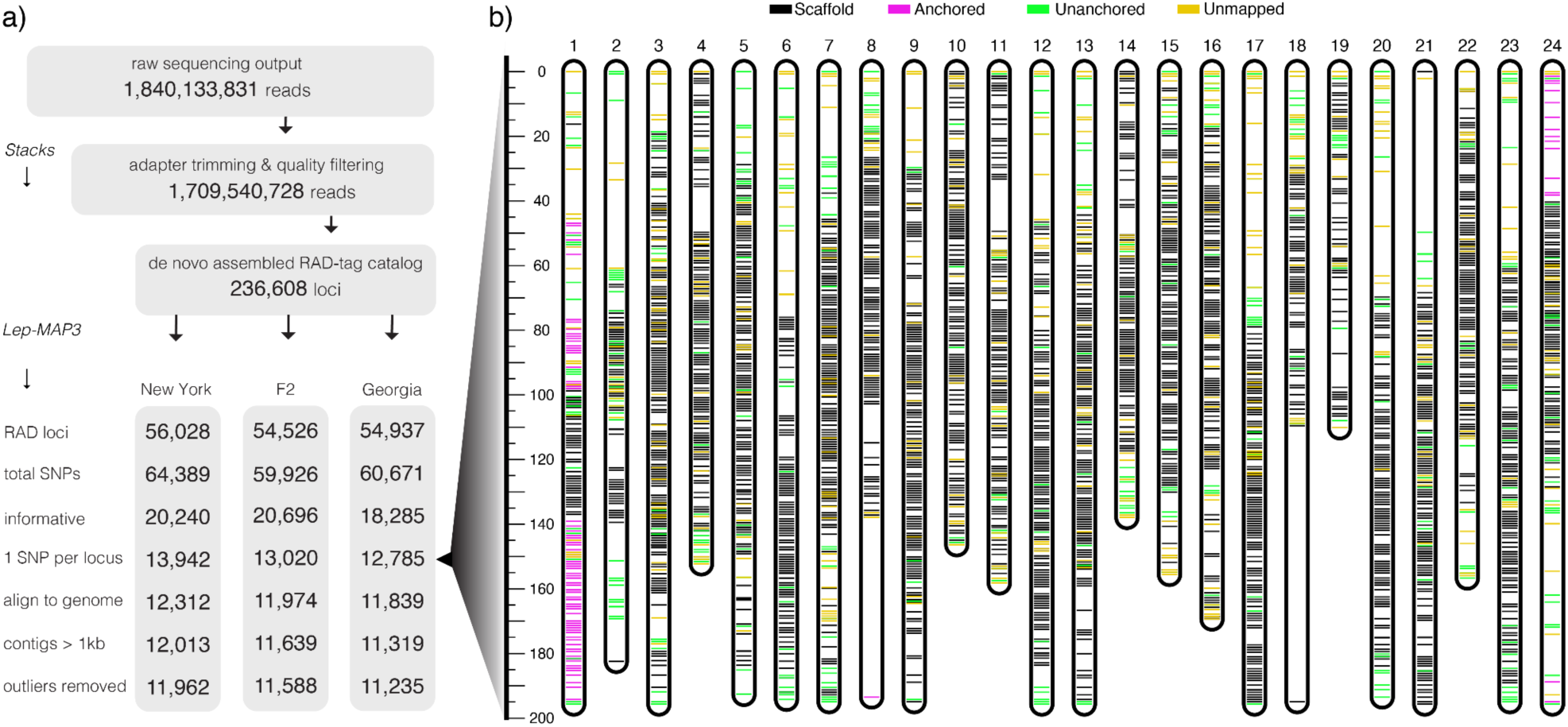
Markers in Female Linkage Maps a) Number of markers in the female linkage maps after each filtering step. b) All SNPs in the Georgia linkage map colored based on their mapping to the linkage-map anchored genome assembly. The y-axis shows genetic distance in centiMorgans. In each linkage group, horizontal black lines represent markers mapping to the main scaffolds in the original assembly, magenta lines are markers added in the anchored assembly, green lines are markers mapping to an unanchored scaffold, and yellow lines are markers that do not map to the reference.

We anchored the three unplaced long scaffolds (> 1 Mb) to the linkage-map guided assembly, adding 15.4 Mb of sequence to the chromosome assembly. Two of these scaffolds, encompassing 7.8 and 4.7 Mb, were added to the beginning of chromosome 1, and the third, encompassing 2.8 Mb, was added to the beginning of chromosome 24. While Lep-Anchor identified an additional 660 scaffolds totaling 15.5 Mb to be anchored to different chromosomes, the resulting Marey maps reveal that these markers are always anchored to the beginning physical positions of each chromosome, despite a large range of genetic positions that place these markers throughout the linkage group (Figure S2). This discrepancy can be attributed to various sources, including misassembly, misrepresentation of repeats, and/or errors in linkage mapping and anchoring. Because further investigation is required to validate the positions of these smaller contigs, we did not include them in our linkage-map guided assembly.

### Synteny analysis in female maps

Comparison of the female linkage maps to the improved reference genome reveals chromosomal rearrangements in all three maps (Figure 5). Our reference genome was assembled from an individual from Georgia, and while the Georgia female linkage map reveals high levels of collinearity with the reference genome sequence as expected, we also see evidence of inversions (reversal of marker ordering in the linkage map compared to the physical sequence). We detected five inversions in the Georgia female linkage map: 0.4 Mb at the beginning of chromosome 1, 1.5 Mb at the end of chromosome 5, 0.9 Mb toward the beginning of chromosome 10, 1.4 Mb toward the beginning of chromosome 12, and 2.1 Mb at the beginning of chromosome 19 (Figure 5). Four of these five inversions (on chromosomes 1, 5, 10, and 12) also appear in the New York and F2 family maps. Additionally, a striking pattern of complete recombination suppression across a wide region (flatlining in genetic distance across > 10 Mb of the physical genome sequence) is seen on chromosomes 6 and 19 of the Georgia female linkage map. These regions may be the signature of inversions that segregate in the Georgia population for which the sampled female was heterozygous.

**Figure 5.**
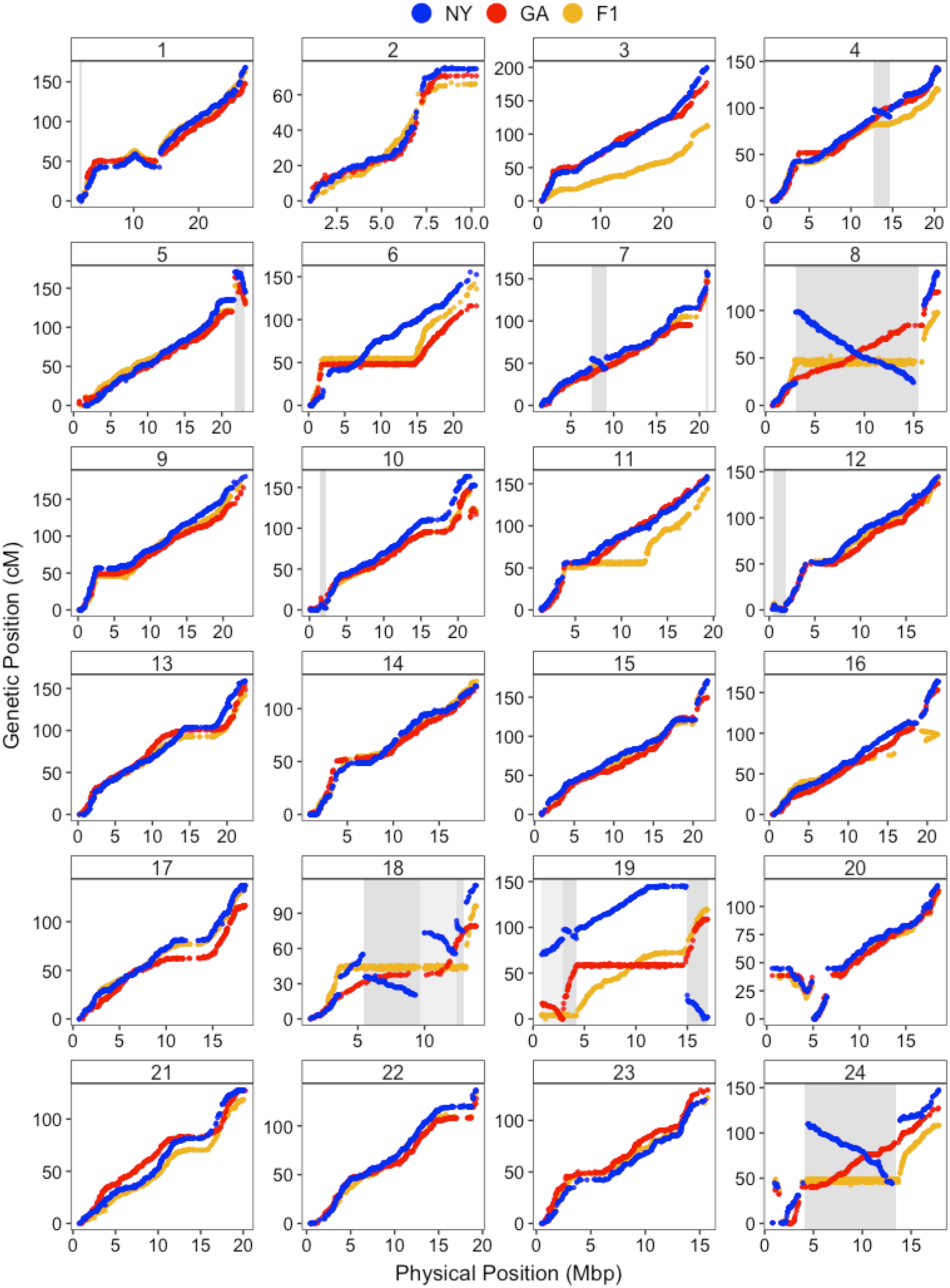
Female Marey Maps Genetic distance (cM) along the physical distance (Mb) of each chromosome is shown for all three females. Each point is a SNP and the marker order in the GA female is shown in red, NY in blue, and their resulting hybrid in yellow. Shaded regions highlight inversions with alternate arrangements in the GA and NY female.

When comparing the New York female linkage map to the Georgia reference genome sequence, we detected a total of 15 chromosomal inversions across 11 of the 24 chromosomes. The inversions range in size from 0.4 to 12.5 Mb, with the largest spanning much of the length of chromosome 8. The majority of chromosomes 18 and 24 are also inverted, with the former having three adjacent inversions at positions 5.5-9.7 Mb, 9.7-12.4 Mb, and 12.4-13.0 Mb, and the latter having the second largest inversion that captures 9.3 Mb (Figure 5). Smaller inversions are seen on chromosome 1 (at position 1.6-2 Mb), chromosome 4 (at position 12.7-14.7 Mb), chromosome 7 (at position 7.4-9.1 Mb), and chromosome 19 (at position 2.9-4.2 Mb). In all, these rearrangements span 44.1 Mb, or 9.2% of the 481.2 Mb chromosome assembly (i.e., the 24 largest scaffolds of the genome).

The F2 family map reveals the effect of these inversions on the recombination landscape in crosses between New York and Georgia (because it reflects meiotic recombination in an F1 daughter with a wild-caught parent from each of these populations). As expected, chromosomal regions with opposite orientations of inversions between these two populations do not recombine in the heterozygous offspring, as revealed by the flatlining of genetic map distances in those regions (Figure 5, yellow data points). Chromosomes 8, 18, and 24, which were previously identified as harboring highly divergent haplotypes in the two studied populations (Lou *et al*. 2018; Wilder *et al*. 2020), show large blocks of suppressed recombination in the hybrid mother of the F2 family map as a result of the inversions. In chromosome 18, recombination is suppressed an additional 1.8 Mb beyond the inversions identified (at position 3.6-5.5 Mb, Figure 5).

Recombination was also suppressed on chromosome 11 in the hybrid female map without evidence of an inversion between the two parental maps in this position. Chromosome 11 was, however, also previously identified as having a large block of SNPs in tight LD and nearly fixed for opposite alleles across the range, supporting the presence of an inversion in this genomic location. To note, highly divergent northern haplotypes associated with this inversion were most common in locations further north (Gulf of Maine and Gulf of Saint Lawrence) of the populations sampled in this study (Lou *et al*. 2018; Wilder *et al*. 2020). While the southern haplotype on chromosome 11 is predominant in both Georgia and New York, the northern haplotype is present in low frequency in New York. Thus, the suppression of recombination in this region of chromosome 11 in the F2 map may be the signature of an inversion that segregates in the New York population, but did not show up in our F1 New York map because the female used to establish the New York map (F1) carried the southern arrangement (collinear with the assembly). A northern (inverted) haplotype was likely introduced by the New York male that became the grandfather of our F2 offspring (see Figure 1), explaining how the F2 offspring became heterozygous for this region. In a similar vein, the recombination suppression seen on chromosome 6 in the Georgia female linkage map is also seen on the F2 family map, while the suppression on chromosome 19 is not, again likely reflecting signatures of inversions that segregate within populations. These four regions of suppressed recombination on Chromosomes 6, 11, 18, and 19 representing putative inversion heterokaryotypes span an additional 33.9 Mb of the chromosome assembly.

### Estimation of recombination rates

As evident from the Marey maps in Figure 5, estimated female recombination rates vary across the genome (Figure 6). We observe increased rates of recombination near the ends of many (but not all) chromosomes and reduced recombination towards the centers, often with drops to near zero, in what are likely centromere regions. While there is a significant negative relationship (R = −0.28, p = 0.016) between chromosome size and average recombination rate in all three maps analyzed together, this trend is non-significant when analyzed for each map separately (Figure 7a). Recombination rates vary among maps (F = 9.26, df = 2,68, p < 0.001), with significantly higher mean recombination rates in New York (7.37 cm/Mb) compared to the Georgia (6.69 cm/Mb) and hybrid F1 (6.51 cm/Mb) female maps, but no significant difference between the latter two (Figure 7a). Chromosomes with and without inversions show no difference in average recombination rate (z = −2.160, p = 0.3761). Variation in fine-scale recombination rates (from the Loess model) is evident across terminal, central, and inverted regions of chromosomes with and without inversions (Figure 7b). ANOVA with Satterthwaite’s method revealed significant differences in recombination rates related to population (F = 8.48, df = 2, p = 0.01), chromosomal region (F = 23.82, df = 3, p < 0.001), and their interaction (F = 66.79, df = 6, p < 0.001). Recombination rates are higher in the terminal ends of all chromosomes, regardless of the presence of inversions. Inverted regions, however, have lower recombination rates compared to regions outside inversions in chromosomes with inversions as well as compared to central regions in chromosomes without inversions. While this pattern is primarily driven by the reduced recombination in inversions in the hybrid map, when considering only the F1 family maps, recombination rates inside inversions are still lower than regions outside inversions (z = 8.70, p < 0.0001) but no different than the central regions of chromosomes without inversions (z = 1.110, p = 0.5079).

**Figure 6.**
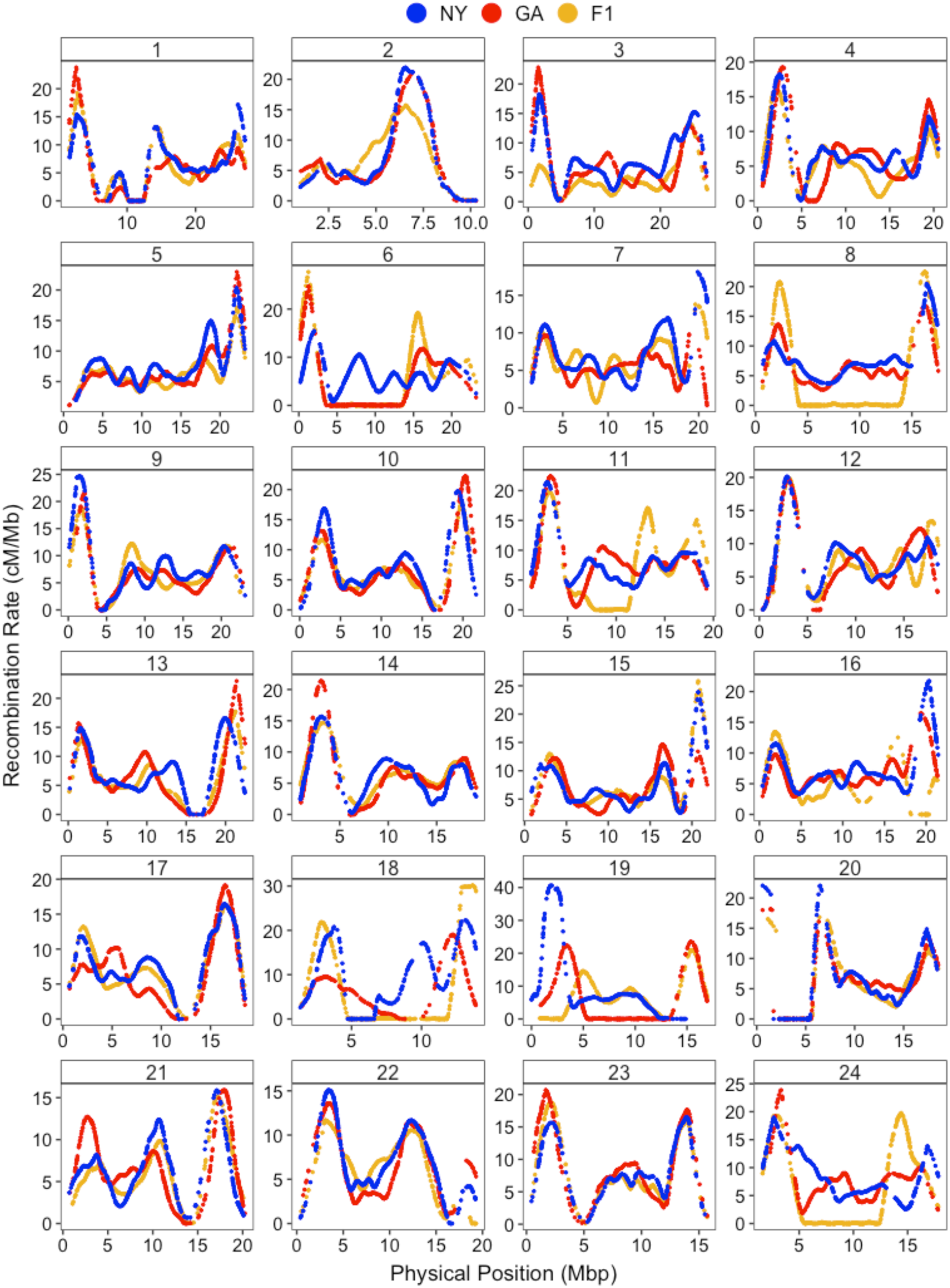
Female Recombination Maps Fitted splines represent the variation in recombination rate as a function of physical distance for the three mapping families. The maps show that most chromosomes have a region (presumably the centromere) where recombination is close to zero in all females and that this region tends to be offset from the center of the chromosome.

**Figure 7.**
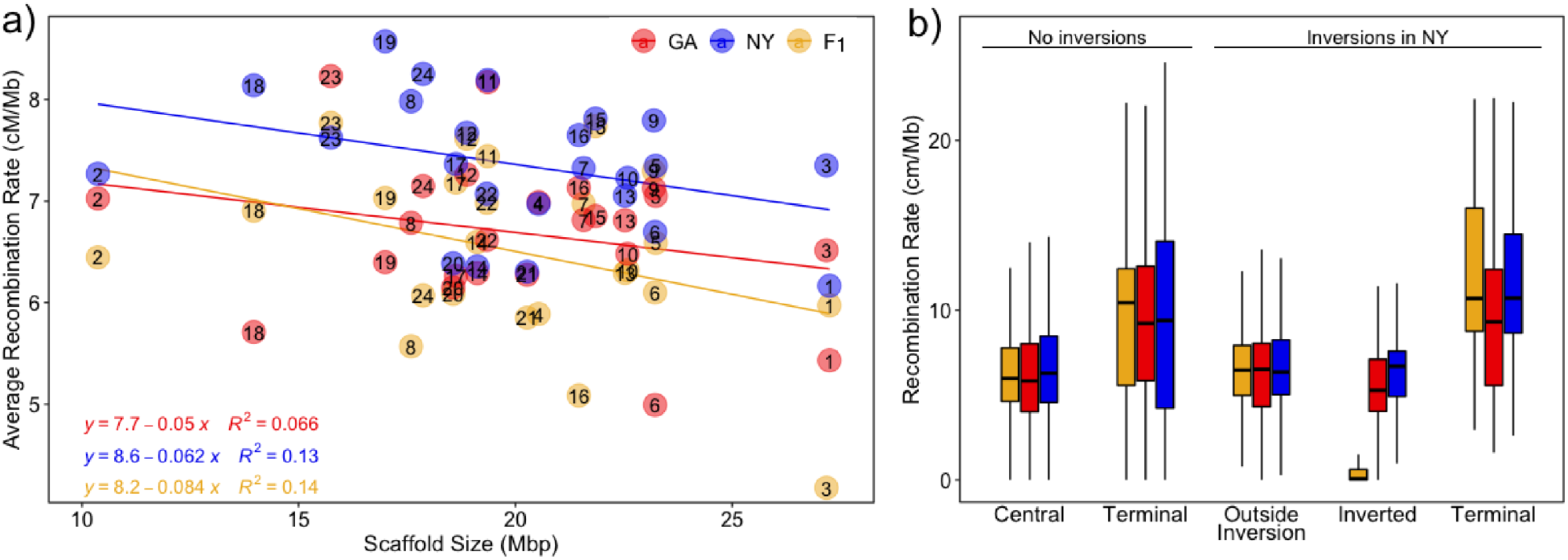
Comparing Recombination Rates a) A comparison across all chromosomes reveals a tendency for higher averaged recombination rates in smaller chromosomes in all three maps, and overall significantly higher recombination rates across chromosomes in NY. b) Across the maps, recombination rates are higher in the terminal ends (10% of each end) of all chromosomes, both those with and without inversions. As expected, the inversions show no recombination in the F1 female that was heterozygous for those regions (yellow), but the inverted regions also have lower recombination rates in both homozygotes (GA and NY shown in blue and red) compared to regions outside inversions.

## Discussion

By building and comparing multiple high-density linkage maps, we found remarkable variation in recombination rates between both sexes and across adaptively divergent populations of the Atlantic silverside. We also validated standing variation in large-scale chromosomal inversions and demonstrated how these inversions suppress recombination in heterozygous individuals.

### Suppressed recombination in males

We showed that the recombination landscape in the Atlantic silverside varies substantially both within and across chromosomes, and between sexes and populations, a pattern that is consistent with other study systems (Kong *et al*. 2010; Smukowski and Noor 2011; Sardell and Kirkpatrick 2020). Males showed virtually no recombination across central portions of all chromosomes (Figure 2). The restriction of recombination to telomeric regions in males has also been demonstrated in other species with female-biased heterochiasmy, which is more common than homoschiasmy in animals (Brandvain and Coop 2012; Sardell and Kirkpatrick 2020). We found that the two F1 family female maps are on average 2.75 times longer than the male maps, one of the most sex-biased recombination rates known for fishes (Figure 3).

A recent metanalysis compared sex-specific recombination rates in 61 fish species, concluding that sex differences in recombination rate are evolutionary labile, with frequent shifts in the direction and magnitude of heterochiasmy that cannot be explained by neutral processes or biological sex differences in meiosis (Cooney *et al*. 2021). Alternative hypotheses include the Haldane-Huxley hypothesis, which posits that recombination may be adaptively suppressed to varying degrees across the genome in the heterogametic sex, in order to prevent X-Y or Z-W crossing over (Haldane 1922; Huxley 1928). However, this probably does not apply to silversides, which exhibit partial environmental sex determination (Conover and Kynard 1981; Duffy *et al*. 2015) and do not appear to have heteromorphic sex chromosomes. In addition, there is no significant correlation between sex determination mechanism and sex-bias in recombination rate across fish species (Cooney *et al*. 2021). Other hypotheses relate to sexual selection and sexual conflict, predicting that patterns of heterochiasmy are a result of stronger selection experienced by one sex or their gametes (haploid selection), but the data to test this are currently lacking for most fish species (Cooney *et al*. 2021). In silversides, partial sexual size dimorphism has been previously documented, with slower growing males experiencing higher size-selective mortality compared to females (Pringle and Baumann 2019). While this is in line with predictions of the sexual conflict hypothesis, which favors suppressed recombination in the sex subject to stronger selection (Sardell and Kirkpatrick 2020), further investigation is warranted to characterize the relationship between sexual conflict and sex-biased recombination rates in silversides.

### Recombination landscapes in females

In the female maps, we found a weak negative correlation between recombination rates and chromosome size, a pattern that is common but not ubiquitous among other species (Stapley *et al*. 2017). As genome size predicts variation in chromosome size (Li *et al*. 2011) and chromosomes of different sizes tend to experience different recombination rates (Haenel *et al*. 2018), a weak negative correlation is expected based on the relatively small genome size of the Atlantic silverside (Tigano *et al*. 2021b). We also discerned differences in fine-scale recombination rates along the genome (Figure 6 and 7), including elevated recombination at the terminal ends of chromosomes, and suppressed recombination in central regions, consistent with patterns in a large variety of taxa (Haenel *et al*. 2018; Peñalba and Wolf 2020).

Looking across populations, we found a tendency for higher average recombination rates in the New York female map compared to both the Georgia and hybrid maps (Figure 7a). Variation in recombination rates among individuals and populations is well-established. Recent work examining *Drosophila* populations demonstrated that natural selection can shape interpopulation differences in recombination rate (Samuk *et al*. 2020). In theory, increased rates of recombination are favored in temporally fluctuating environments when fitness optima change rapidly (Charlesworth 1976), while lower recombination rates are favored when adaptive combinations of alleles are at risk of dissociation by maladaptive gene flow (Kirkpatrick and Barton 2006). In the case of silversides, temporal fluctuation in the environment is more pronounced in the north (Conover and Kynard 1981; Duffy *et al*. 2015), while higher levels of gene flow disproportionately affect southern populations (Lou *et al*. 2018; Wilder *et al*. 2020). Across the latitudinal range of Atlantic silversides, the length of the growing season abruptly shifts around 38°N (Conover and Kynard 1981; Duffy *et al*. 2015), which is just 2° south of our northern sampling site in New York. Thus, while the two focal populations in our study gave us a glimpse into the variation in recombination across populations, a thorough investigation of recombination patterns across the species range is needed to determine the geographic distribution of this variation.

### Chromosomal inversions

We detected a total of 15 chromosomal inversions across 11 of the 24 chromosomes. These inversions range in size from 0.4 to 12.5 Mb and in total span 44.1 Mb or 9.2% of the silverside genome. We detected four additional putative inversions, presumably heterozygous in the Georgia and hybrid maps, ranging in size from 1.8 to 12.9 Mb and together span an additional 33.9 Mb or 7% of the silverside genome. Overall, these rearrangements span 76.6 Mb, or 15.9% of the 481.2 Mb chromosome assembly. These findings add to a growing body of studies that implicate inversions as important drivers of evolutionary change. A powerful mechanism for protecting co-adapted alleles from dissociation, large inversions are widespread and typically span many genes: a recent review showed that the average reported inversion size in both plants and animals is 8.4 Mb, ranging from 130 kb to 100 Mb, and contain an average of 418 genes (Wellenreuther and Bernatchez 2018). Here, we identified a subset of the 662 inversions affecting 23% of the genome recently reported to be segregating between southern and northern populations of Atlantic silversides, inferred from alignment of independent genome assemblies (Tigano *et al*. 2021a) (Figure 8). Despite only identifying 19 inversions, which overlap 62 of the 662 previously identified by Tigano *et al*. (2021a), the inversions we identified affect nearly 16% of the genome. Our study, which provides independent evidence that clearly confirms the presence and impact of the larger inversions detected in the genome, certainly underestimates the total number of rearrangements in the silverside genome. This is a reflection of the ascertainment bias of reduced genome representation methods (such as the RAD genotyping used here), which only have the resolution to detect relatively large inversions. In addition, linkage mapping can be biased by the individuals used to establish a pedigree; a single pedigree cannot fully capture the true recombination landscape of the focal population, and we only observe inversions segregating in our specific founding individuals. Moreover, this method only considers recombination events in gametes that resulted in offspring and does not characterize recombination in unsuccessful gametes. However, our pedigree-based approach provides a direct estimate of genetic linkage by observing the inheritance of alleles in a few families, allowing us to robustly distinguish recombination rates among individuals of the parental generation, including the different sexes and populations. Compared to population-based inferences (for estimating recombination and detecting inversions), genetic maps are affected to a much lesser extent by demography and selection acting across evolutionary times and provide a key resource for future comparative genomic and QTL studies in this species (Sarropoulou and Fernandes 2011; Samuk and Noor 2021).

**Figure 8.**
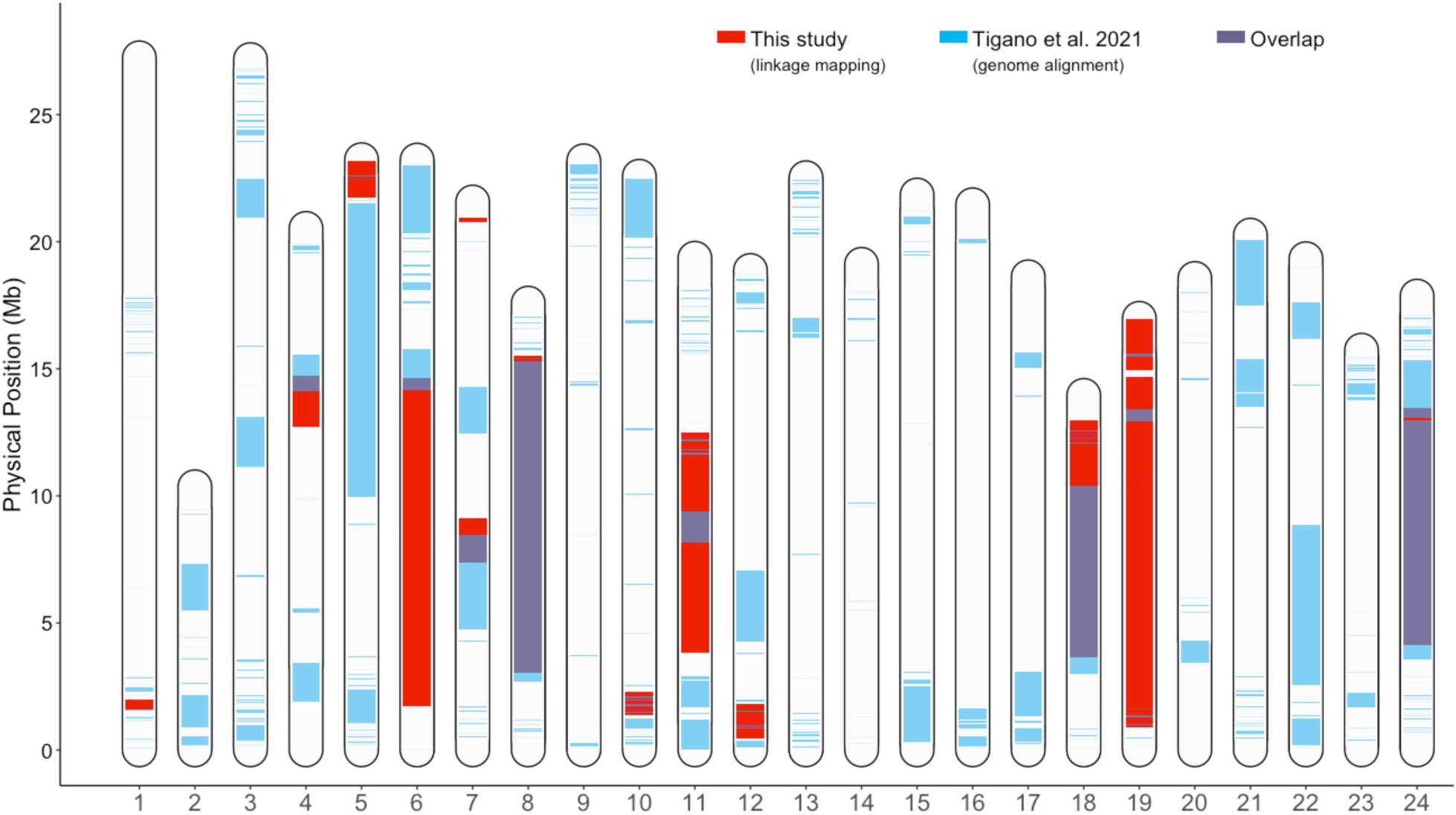
Inversions Distribution of inversions identified in our study, compared to previous reports (Tigano et al. 2021).

### Genomic patterns associated with adaptive divergence

Structural variation within the genome can promote genomic divergence by locally altering recombination rates. The key evolutionary effect of inversions is that they suppress recombination in a heterozygous state (Sturtevant and Beadle 1936). This study demonstrated that inversion polymorphisms between locally adapted Atlantic silverside populations suppress recombination in inter-population hybrids. Suppressing recombination is an efficient way to preserve linkage between favorable combinations of locally adapted alleles but is advantageous only when populations experience gene flow (Faria *et al*. 2019b). Populations of the Atlantic silverside south of Cape Cod (including both Georgia and New York) show high connectivity across this broad geographic range that spans the steep latitudinal temperature gradient of the North American Atlantic coast (Lou *et al*. 2018; Wilder *et al*. 2020). Hatching in the intertidal zone in the spring, silversides move up to 170-km offshore to overwinter (Conover and Murawski 1982), and extensive mixing between spawning sites has been documented (Clarke *et al*. 2009; Wilder *et al*. 2020). The discovery of chromosomal rearrangements that suppress recombination between populations suggests a possible mechanism that could preserve the association between locally favorable alleles and as such help maintain combinations of locally adaptive traits (Therkildsen *et al*. 2019).

Inversions can capture alleles that control adaptive traits into a single complex block to prevent their dissociation by reducing recombination in heterokaryotypes, while the majority of the genome is homogenized by gene flow. Although we saw no recombination within the large inversions among the 221 offspring examined here, the recombination reduction in heterokaryotypes, however, is not necessarily complete on a population scale because viable recombinant gametes may arise by double crossing over or by gene conversion (Sturtevant and Beadle 1936; Chovnick 1973), but this tends to occur at low rates. Furthermore, inversion polymorphisms are not static, but continue to evolve after establishment. Inversion dynamics are thus complex and depend on the relative roles of selection, drift, mutation, and recombination, all of which change over time and have implications for the inversion itself and the evolution of the populations (Faria *et al*. 2019b).

An outstanding question regarding the role of inversions in adaptive evolution is whether they become targets of strong selection because of their content or because they generate mutations or gene disruptions at breakpoints (Kirkpatrick 2010; Wellenreuther and Bernatchez 2018). Previous work has shown that linked genes in the major inversion regions are enriched for functions related to multiple local adaptations in silversides: markers on outlier sections of chromosome 8, 18, and 24 were enriched for gene ontology terms related to polysaccharide metabolic processes, meiotic cell cycle, cartilage morphogenesis, regulation of behavior, and regulation of lipid storage, and these functions all relate to traits that show adaptive divergence in this species (Wilder *et al*. 2020). This functional enrichment could suggest that gene content may play an important role in the origin and maintenance of inversions and could indicate that the inversions may act as supergenes to maintain coinheritance of adaptive alleles. An important aspect of supergenes is that they allow switching between discrete complex phenotypes and can maintain stable local polymorphism without the generation of maladaptive intermediates (Thompson and Jiggins 2014). While we have evidence from our linkage maps that some structural variants are polymorphic within populations (e.g. inversions on chromosome 6 and 19 in Georgia and on chromosome 11 in New York), determining whether inversions are indeed acting as supergenes requires further work to disentangle phenotype-genotype association and examine their frequencies within and among populations, as well as to rule out alternative hypotheses (e.g., inversions disrupt associations of gene-regulatory elements).

## Acknowledgements

We thank Harmony Borchardt-Wier for help with DNA extractions, Steve Bogdanowitz for help with preparation of the RAD-seq libraries, Pasi Rastas for discussions and advice on implementing and interpreting Lep-Map3 and Lep-Anchor analyses, and members of the Therkildsen lab for their thoughtful feedback on the manuscript. This study was funded through a National Science Foundation grant to NOT (OCE-1756316) and OCE-1756751 to HB.

## Data Accessibility

Raw data from the RADseq libraries will be available under NCBI BioProject accession number PRJNA771889. Scripts for all analyses will be available at http://github.com/therkildsen-lab/silverside-linkage-maps

